# Mutations in a barley cytochrome P450 gene enhances pathogen induced programmed cell death and cutin layer instability

**DOI:** 10.1101/2021.03.09.434546

**Authors:** G. Ameen, S. Solanki, L. Sager-Bittara, J. Richards, P. Tamang, T.L. Friesen, R.S. Brueggeman

**Author notes:** Corresponding Author: Robert S. Brueggeman.

## Abstract

Disease lesion mimic mutants (DLMMs) are characterized by spontaneous development of necrotic spots with various phenotypes designated as necrotic (*nec*) mutants in barley. The *nec* mutants were traditionally considered to have aberrant regulation of programmed cell death (PCD) pathways, which have roles in plant immunity and development. Most barley *nec3* mutants express cream to orange necrotic lesions contrasting them from typical spontaneous DLMMs that develop dark pigmented lesions indicative of serotonin/phenolics deposition. Also, barley *nec3* mutants grown under sterile conditions did not exhibit necrotic phenotypes until inoculated with adapted pathogens suggesting that they are not typical DLMMs. The F_2_ progeny of a cross between *nec3-*γ1 and variety Quest segregated as a single recessive gene post inoculation with *Bipolaris sorokiniana*, the causal agent of the disease spot blotch. *Nec3* was genetically delimited to 0.14 cM representing 16.5 megabases of physical sequence containing 149 annotated high confidence genes. RNAseq and comparative analysis of wild type and five independent *nec3* mutants identified a single candidate cytochrome P450 gene (HORVU.MOREX.r2.6HG0460850) that was validated as *nec3* by independent mutations that result in predicted nonfunctional proteins. Histology studies determined that *nec3* mutants had an unstable cutin layer that disrupted normal *Bipolaris sorokiniana* germ tube development.

**AUTHOR SUMMARY:** At the site of pathogen infection, plant defense mechanisms rely on controlled programmed cell death (PCD) to sequester biotrophic pathogens that require living cells to extract nutrient from the host. However, these defense mechanisms are hijacked by necrotrophic plant pathogens that purposefully induce PCD mechanism to feed from the dead cells facilitating further disease development. Thus, understanding PCD responses is important for resistance to both classes of pathogens. We characterized five independent disease lesion mimic mutants of barley designated necrotic 3 (*nec3*) that show aberrant regulation of PCD responses upon pathogen challenge. A cytochrome P450 gene was identified as *Nec3* encoding a Tryptamine 5-Hydroxylase that functions as a terminal serotonin biosynthetic enzyme in the Tryptophan pathway of plants. The *nec3* mutants have disrupted serotonin biosynthesis resulting in expansive PCD, necrotrophic pathogen susceptibility and cutin layer instability. The *nec3* mutants lacking serotonin deposition in pathogen induced necrotic lesions show expansive PCD and disease susceptibility suggesting a role of serotonin to sequester PCD and suppress pathogen colonization. The identification of *Nec3* will facilitate functional analysis to elucidate the role serotonin plays in the elicitation or suppression of PCD immunity responses to diverse pathogens and effects it has on cutin layer biosynthesis.

## INTRODUCTION

Programmed cell death (PCD) is a tightly regulated physiological response of plant and animal cells that functions in development, cell differentiation, cell number homeostasis, and immunity (1). In plants, PCD is activated by environmental cues that include biotic stress induced by pathogens. Specialized or adapted plant pathogens evolved to produce and utilize virulence effectors to facilitate host penetration and colonization by manipulating host cellular machinery to circumvent basal resistance mechanisms and induce inappropriate physiological responses that promote access to nutrients and life cycle completion (2). The selective pressures exerted by adapted pathogens led to the counter-evolution of a complex innate immune systems to recognize the diverse repertoire of pathogen virulence effectors (3–6) or more commonly their manipulation of targeted host proteins (7). This direct or indirect recognition of ‘non-self’ typically induces effector triggered immunity (ETI), which is characterized by a strong PCD response known as the hypersensitive response (HR). The HR functions to sequester and inhibit further colonization by biotrophic pathogens. Although the scientific community is beginning to elucidate these mechanisms in plants, disease lesion mimic mutants (DLMMs) that exhibit spontaneous and induced PCD responses are an underutilized tool that could be studied to fill gaps in understanding how PCD responses function in plant immunity.

The plant innate immune system was categorized into distinct layers. The first line of defense is known as <u>p</u>athogen <u>a</u>ssociated <u>m</u>olecular <u>p</u>atterns (<u>PAMP</u>) <u>t</u>riggered immunity (PTI). Many of the characterized PTI mechanisms were designated as “basal” or “nonhost” defense responses because they are triggered at the cell surface early in the host-parasite interaction by detection of microbial molecules conserved across genera or species that are indispensable for fitness. Thus, PTI provides resistance against a broad taxon of microbes or potential pathogens. The PTI responses include callose deposition at the point of pathogen penetration, induced expression of pathogenesis-related (PR) genes, a quick transient reactive oxygen species (ROS) burst, and in some responses, PCD at the site of attempted pathogen penetration. This PCD viewed as necrotic lesions is typically accompanied by the deposition of phenolic compounds resulting in dark pigmentation (8). PTI is induced by transmembrane cell surface receptors known as <u>p</u>attern <u>r</u>ecognition <u>r</u>eceptors (PRRs) that function in heterologous complexes (9) that activate underlying cytosolic signaling cascades including the mitogen activated protein kinase (MAPK) pathways (10). The PRRs typically contain diverse extracellular receptor domains, a transmembrane domain and are grouped into two classes, 1) those that contain an intracellular kinase signaling domain known as receptor-like kinases (RLKs), or 2) those that are missing the kinase domain known as receptor-like proteins (RLPs) (8).

The PRR complexes identify PAMPs, which are microbial molecules conserved across genera or species that are indispensable for fitness. Bacterial PAMPs include elongation factor EF-Tu (11), lipopolysaccharides (12) and the flagellin subunits flg22. Flagella are required for motility by many gram-negative bacteria (13). In *Arabidopsis thaliana*, the RLK FLS2 directly interacts with flg22 to elicit PTI defense responses. Known fungal PAMPs are chitin, (14), β-glucan (15) and ergosterol (16). The detection of fungal derived chitin fragment in rice requires two PRRs that contain extracellular LysM domains, the LysM RLK OsCERK1 and the LysM RLP CEBiP, to elicit chitin-mediated PTI signaling (17). PTI-like responses are also elicited by <u>d</u>amage <u>a</u>ssociated <u>m</u>olecular <u>p</u>atterns (DAMPs) known as DAMP triggered Immunity (DTI). The DAMPs are host derived extracellular matrix subunits such as oligogalacturonide residues (OGs) released from plant cell walls upon partial degradation by pathogens. The detection of OGs by wall associated kinase (WAK) PRRs sound the alarm of compromised cell integrity, indicating pathogen ingress or challenge eliciting the DTI defense responses (18,19).

The selective pressures exerted by pathogens forced plant innate immune systems to counter-evolve to recognize pathogen virulence effectors (3–6) or more commonly their manipulation of targeted host proteins (7). This has been well characterized with *Psuedomonas syringae* effectors that inhibit FLS2 PRR-mediated signaling following flg22 perception (20). However, the zig-zag model postulates that during the host-parasite molecular arms race (21) plants counter-evolved cytoplasmically localized immunity receptors, typically with nucleotide binding site-leucine rich repeat (NLR) protein domain architecture, to recognize the presence of these effectors. NLR activation elicits ETI resulting in a higher amplitude HR, which is considered the second layer of plant defenses.

Once a virulence effector is recognized by a cognate NLR immunity receptor, it effectively becomes a pathogen avirulence protein in the presence of the cognate resistance gene. For biotrophic pathogens that require living host cells to feed, these effectors no longer facilitate colonization but rather alert the host to their presence, eliciting PCD that kills the cells they are feeding from, effectively stopping the colonization process. Thus, HR is critical to plant innate immunity against biotrophic plant pathogens including viruses, bacteria, fungi, oomycetes, and invertebrates (22). However, the necrotrophic pathogens such as *Prastagonospora nodorum* (23) and *Pyrenophora teres* (24) have adapted to hijack these gene-for-gene immunity mechanisms by evolving necrotrophic effectors (NEs) that purposely alert the host immune system of their presence through immunity receptor activation. These inverse gene-for-gene interactions (25) initiate PCD responses, which the necrotrophic pathogens utilize to facilitate disease formation because they acquire nutrient from the resulting dying and dead tissues. Thus, necrotrophic pathogens can complete their lifecycle on the host by facilitating further disease development through necrotrophic effector triggered susceptibility (NETS) (24). Both biotrophic and necrotrophic pathogens elicit PCD immunity responses in plants with different outcomes, incompatibility (resistance) –vs-compatibility (susceptibility), respectively. Thus, this outcome is determined by the lifestyle of the pathogen and the timing of the responses. Knowledge of PCD pathways is important to understand resistance and susceptibility mechanisms in crop plants when interacting with both classes of pathogens.

Pathogen induced HR begins with an efflux of hydroxide and potassium into the apoplastic spaces and an influx of calcium and hydrogen ions into the cytoplasm (26,27). This is followed by the production of ROS, which results in an oxidative burst in the cells undergoing the HR that includes super oxide anions, hydrogen peroxide, and hydroxyl radicals (28). Besides ROS, the production of reactive nitrogen species (RNS) and the phytohormone salicylic acid (SA) are also produced (27). The species and amount of ROS molecules present and their compartmentalization have been implicated in HR signaling (29,30).

Phenolic secondary metabolite compounds including serotonin are synthesized and deposited in cells at the site of PCD and surrounding the necrotic lesions that sequester or harbor biotrophic and necrotrophic pathogens during immunity responses (31). Serotonin acts as a potent antioxidant by scavenging reactive oxygen species (ROS) and is incorporated into the cell wall at infection sites. Thus, serotonin can function to strengthen host cell-walls and inhibit pathogen growth to sequester pathogens in the HR lesion (31). Secondary metabolites including serotonin may also play a role to sequester PCD lesion expansion. Thus, optimizing the plants immune response to effectively stop the pathogen in a regulated foci of dead cells yet retaining the photosynthetic capability of the surrounding leaf tissue (32,33).

The Sekiguchi lesion (*sl*) mutant of rice produces tan to orange necrotic lesions similar in appearance to the barley *nec3* mutants characterized in this study. It has been shown that the *sl* mutant is incapable of serotonin biosynthesis and lacks the dark phenolic deposition in its necrotic lesions due to the missing serotonin deposition (31). It was shown that exogenous addition of serotonin to the *sl* mutant restored the dark deposition of the phenolic compound in its induced lesions and increased resistance to the necrotrophic pathogen *Bipolaris oryzae* the causal agent of the rice brown spot disease (31). Additionally, the addition of serotonin suppressed the fungal hyphal growth in culture media. Interestingly, we utilized five independent *nec3* mutants in this study and four of them resulted in pathogen induced PCD responses lacking the dark phenolics typical of pathogen induced lesions that resembled the rice *sl* mutant lesions. However, one of the allelic *nec3* mutants (*nec3.d*) produced the dark phenolics typically observed in necrotrophic pathogen induced lesions and DLMMs.

Although radiation and chemical induced mutant populations in crop plants have been available for close to 100 years (34), the advancement of molecular techniques are now allowing for the identification and functional analysis of genes underlying DLMMs (35–38). The identification of genes underlying DLMM phenotypes by forward and reverse genetics approaches will help answer questions concerning unique and differentially regulated PCD pathways that are important in abiotic and biotic stresses, and PCD processes involved in plant development (36). However, few DLMMs have been thoroughly characterized leaving these collections as underutilized resources.

In barley several DLMMs have been described (36), but only two genes have been identified, *Hvnec1* and *mlo*. The *Hvnec1* gene encodes a cyclic-gated ion channel protein (39,40) with sequence homology to the *Arabidopsis thaliana HLM1* gene (41). Like *HLM1*, *Hvnec1* has increased pathogenesis-related (PR) protein expression and produces spontaneous necrotic lesions and leaf tip necrosis with increased susceptibility to certain pathogens (42,43). The *mlo* gene confers increased resistance to the ascomycete biotrophic fungal pathogen *Erysiphe graminis* f. sp. *hordei*, the causal organism of the disease powdery mildew. The *mlo* gene has been deployed in Northern European barley as a source of durable powdery mildew resistance for 37 years (44). However, in the absence of the disease *mlo* induces an ~4% yield penalty due to its DLMM phenotype which results in loss of photosynthetic potential, thus is only economically effective under high disease pressure (45). A second cost of *mlo* deployment is enhanced susceptibility to several necrotrophic pathogens including *Bipolaris sorokiniana* the causal agent of the barley disease spot blotch (46), *Fusarium graminearum* the cause of fusarium head blight (47), *Ramularia collo-cygni* the causal agent of Ramularia leaf spot (48) and *Magnaporthe oryzae* the causal agent of the rice blast disease (49). The *Mlo* gene encodes a ROP like G-protein that appears to be a suppressor of PCD and is conserved and found in other species including pea, *Arabidopsis* and tomato (50).

In barley, both chemical and irradiation mutagenesis have been utilized to induce a large collection of DLMMs (51). One interesting DLMM mutant designated *nec3* develops distinctive cream to orange necrotic lesions (Fig. 1) and was further characterized and mapped utilizing morphological markers (52) to the centromeric region of barley chromosome 6HS ~29.2 cM distal of the *rob1* (orange lemma 1) locus (51–53) and about 16.7 cM distal of the msg36 (male sterile genetic 36) locus (53). The *nec3* gene was more precisely positioned using SNP markers with *nec3.d* shown to be associated with the SNP markers 1_00616 to 1_0882 (position 70.15 cM) on chromosome 6H using the Bowman backcross-derived line BW629 (37). The *nec3.e* mutant was shown to be associated with SNP markers 1_0427 to 1_0494 (positions 56.64 to 70.15 cM) on chromosome 6H using the Bowman backcross-derived line BW630 (37).

**Figure 1.**
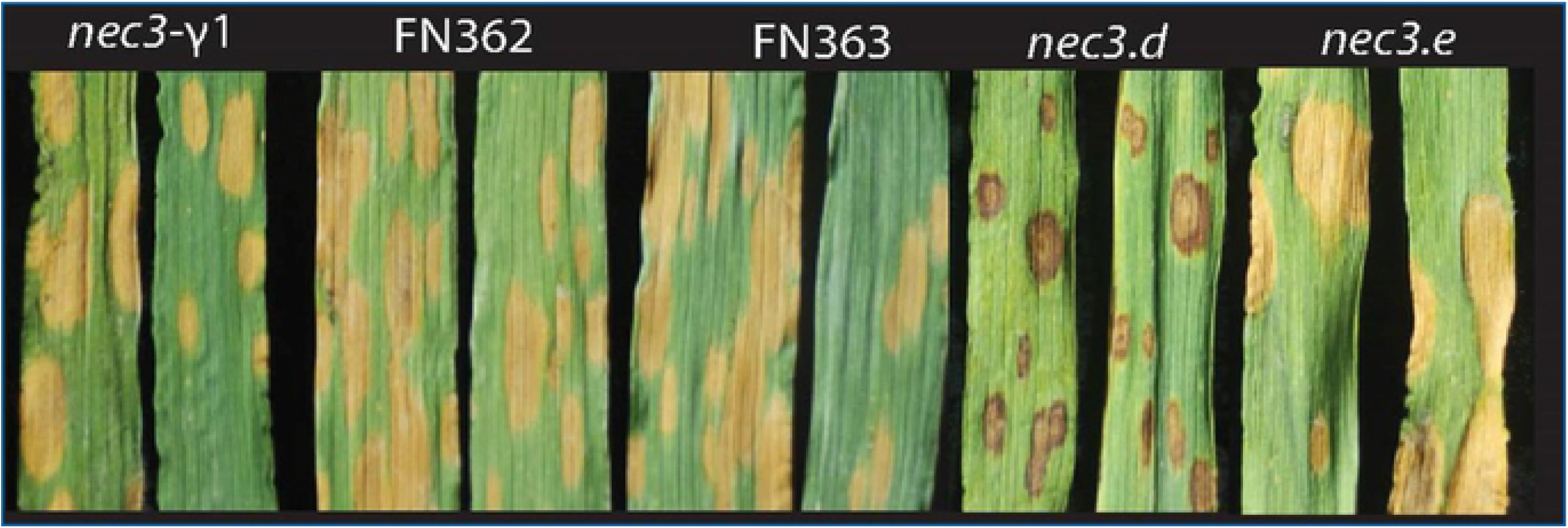
The pathogen induced phenotypic reaction are the orange tan necrotic lesions of the *nec3* mutants. The *nec3* mutants are shown from left to right (*nec3-γ1*, *nec3*.l, *nec3*.m, *nec3*.d and *nec3*.e) after infection with *Bipolaris sorokiniana* isolate ND85F.

Utilizing the Affymetrix Barley1 GeneChip assay, an attempt was previously made to identify the *Nec3* gene, however the results of the study were unsuccessful (54). Despite the gene not being found it was observed that the *nec3.l* and *nec3.m* mutants (Fig. 1) had differential regulation of abiotic stress responsive genes involved in cold and drought stress compared to the wild type. The highest fold change differences in biotic stress related genes were identified as belonging to lipid transfer, pathogen defense and cell wall modifying enzymes (54). Interestingly, mutant seedlings grown for RNA isolation in growth chambers did not express the *nec3* phenotype that was observed on seedlings at the same growth stages consistently expressed under greenhouse conditions. These observations suggested that environmental influences, abiotic or biotic, conditioned the expression of the *nec3* phenotype.

The *nec3* mutants express a distinctive PCD phenotype with tan to orange necrotic lesions lacking the dark phenolics deposition typically associated with DLMMs and pathogen induced necrotic leaf spot diseases. In this study, we report that the *nec3* mutants are not typical spontaneous DLMM, because the necrotic phenotype is only induced by pathogens. This included several species of *Ascomycete* pathogenic fungi, both necrotrophs and biotrophs and the bacterial pathogen *Xanthomonas translucens.* All the pathogens tested that induced the *nec3* phenotype were specialized and penetrate the host by disrupting the cell wall and plasma membrane. We also show that the *nec3* mutants have an unstable cutin layer that peels away from the leaf surface when in contact with *Bipolaris sorokinina* germ tubes. This aberrant interaction alters the pathogen’s developmental signaling resulting in profuse branching of the fungal germ tubes on the mutant leaf surface. In this study, the *nec3* gene was genetically delimited and RNAseq comparative analyses between WT Bowman and the *nec3-*γ1 mutant (Bowman background) identified a Cytochrome P450 gene (HORVU.MOREX.r2.6HG0460850) as a strong candidate *Nec*3 gene. Allele analysis of five independent *nec3* mutants identified predicted deleterious mutations in HORVU.MOREX.r2.6HG0460850 validating it as *Nec3*. Interestingly, *Nec3* is an ortholog of the *sl* gene of rice which catalyzes the conversion of tryptamine to serotonin and the *sl* mutant expresses a similar lesion mimic phenotype as *nec3* (31,55). The identification of the barley *Nec3* gene will facilitate further functional analysis to elucidate the roles it plays in the elicitation or suppression of PCD responses to diverse pathogens and the effects it has on cutin layer biosynthesis.

## RESULTS

### Pathogen Induction of the nec3 Phenotype

The *nec3* symptoms consistently occurred under normal greenhouse conditions yet were not observed when grown in growth chambers (56). To determine if the *nec3* lesions were induced under sterile environmental conditions in the greenhouse or induced by pathogen challenge, as were observed when *nec3-γ1* was inoculated with *B. sorokiniana*, a sterile isolation box experiment was conducted in the greenhouse. The wildtype and mutant plants grown under sterile conditions did not express the *nec3* phenotype and grew to the adult plant stage (Feekes 10.5; full head emergence) without showing any necrotic lesions. The non-inoculated *nec3* seedlings grown on the greenhouse bench outside the isolation box exhibited the *nec3* phenotype at the second leaf stage and continued to develop these distinctive lesions (Figure 1) through the adult plant stages. Histological characterization of the fungal structures that grew from the lesions of the uninoculated *nec3* plants after plating on water agar showed that they were primarily colonized by *Blumeria graminis*, which is endemic in the greenhouses at Washington State University and North Dakota State University where these experiments were conducted.

Inoculations performed under controlled environmental conditions in growth chambers with the most sterile environment possible determined that the typical *nec3* lesions were elicited on *nec3-γ1* mutant seedlings by the necrotrophic ascomycete fungal pathogens *Bipolaris sorokiniana*, *Pyrenophora teres* f. *teres, Pyrenophora teres* f. *maculata*, and *Pyrenophora tritici repentis*, as well as the biotrophic ascomycete fungal pathogen *Blumeria graminis* (Figure 2). Infiltration inoculations with the bacterial pathogen *Xanthomonas translucens* pv *undulosa* also induced the *nec3* phenotype (Figure 2). The *nec3* phenotype was not induced when *nec3-γ1* mutant seedlings were inoculated with the basidiomycete biotrophic fungal pathogen *Puccinia graminis* f. sp. *tritici* race QCCJB that is virulent on cv Bowman which carries the *Rpg1* stem rust resistance gene (Figure 2). The *nec3* phenotype was also not induced when *nec3-γ1* mutant seedlings were inoculated with *P. graminis* f. sp. *tritici* race HKHJC that is avirulent on cv Bowman due to resistance provided by *Rpg1* (Data not shown). The *nec3* phenotype was also not induced by the barley non-host necrotrophic pathogens *Cercospora beticola* and *Parastagonospora nodorum.*

**Figure 2.**
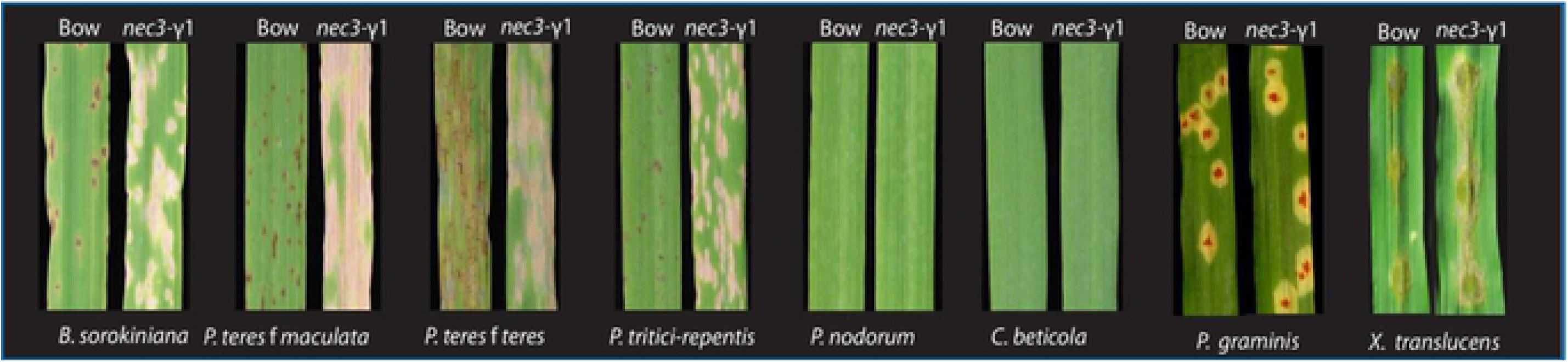
The phenotypic reaction induced by several biotrophic and necrotrophic pathogens on the Bowman wildtype and the *nec3-γ1* mutant. The *nec3* phenotypic reaction was induced by *Bipolaris sorokiniana*, *Pyrenophora teres* f. *teres*, *Pyrenophora teres* f. *maculata, Pyrenophora tritici repentis*, and *Xanthomonas translucens* pv *undulosa*. Each panel shows the typical reaction to the pathogen labelled below with wildtype Bowman (Bow) on the left and the *nec3-γ1* mutant on the right as labeled above. Inoculations with *Puccinia graminis* f. sp. *tritici race QCCJB, Cercospora beticola* and *Parastagonospora nodorum* did not induce the *nec3* phenotype.

### Allelism Crosses

Allelism crosses were completed in the field to determine if the four confirmed *nec3* mutants, *nec3*.d, *nec3*.e, *nec3*.l, and *nec3*.m were allelic to *nec3-γ1*. Crosses between *nec3-γ1* and the mutants, *nec3.d, nec3.e* and *nec3.l e* were successful but due to the head sterility effect of the *nec3* mutants, only 2-5 F_1_ seed were planted from each allelism test cross. 100% of the F_1_ plants displayed the *nec3* phenotype following inoculation with *B. sorokiniana* isolate ND85F under greenhouse conditions (Figure 3). The F_2_ seed of each cross were planted in field rows containing 20 individual F_2_ plants adjacent to a spot blotch disease nursery containing spreader rows inoculated with *B. sorokiniana* isolate ND85F. 100% of the F_2_ individuals exhibited the *nec3* phenotype (Figure 3) demonstrating that the *nec3*-γ1 mutation was allelic to the other known *nec3* mutants. Interestingly, the *nec3.d* mutant which expresses dark lesions typical of phenolics build up that are smaller in size compared to the other *nec3* mutants expressed a range of lesion phenotypes, from the typical *nec3.d* to the *nec3.γ1* phenotype and a blending of both in the F_2_ progeny (Figure 3). This result of the *nec3.γ1* x *nec3.d* crosses suggested that segregation of the two *nec3* alleles had a blending effect on the lesion phenotypes.

**Figure 3.**
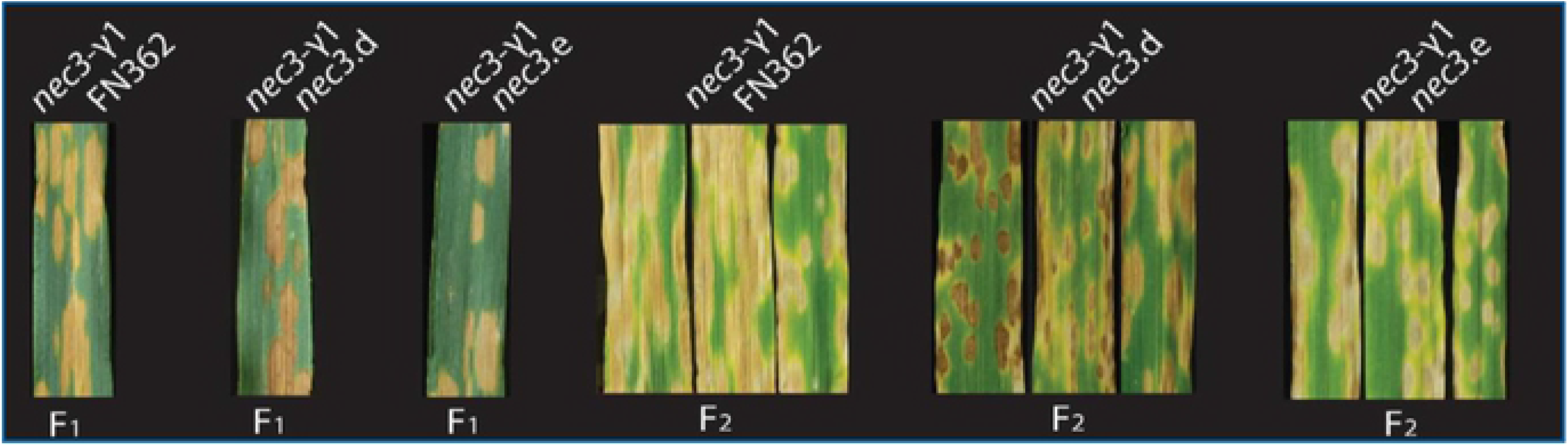
The typical phenotypes of the F_1_ and F_2_ progeny of *nec3* allelism tests. The first three panels show the phenotype of F_1_ progeny of the *nec3* mutants from allelic test crosses after infection with *Bipolaris sorokiniana* isolate ND85F. The progeny from the allelism test crosses were advanced to the F_2_ generation and assayed for the *nec3* phenotype post *B. sorokiniana* isolate ND85F inoculation. The allelism test crosses are shown above and generations shown below.

### Culture Filtrate Infiltration

Infiltrations of WT Bowman and WT Steptoe and the *nec3* mutants *nec3-γ1, nec3.d* and *nec3.e* in the Bowman background and *nec3.l* and *nec3.m* in the Steptoe background showed differential reactions after infiltration with *B. sorokiniana* isolate ND85F culture filtrates and controls infiltrated into secondary leaves (S1 Figure). The infiltrations containing culture filtrates + Fries Media, culture filtrates + MOPS buffer, and culture filtrates + MOPS + pronase on the mutants *nec3-γ1, nec3.d, nec3.e*, *nec3.l* and *nec3.m* showed typical *nec3* PCD as previously described with a lack of defined margins around the lesions (Figure S1). The control infiltrations containing Fries Media + MOPS buffer + pronase did not induce a reaction on the mutants or WT leaves. The WT Bowman leaves showed no response to any of the infiltrations, culture filtrates or control. However, the WT Steptoe seedlings displayed a PCD response to all the infiltrations containing culture filtrates that were different than *nec3.l* and *nec3.m*, the *nec3* mutants in the Steptoe background. Steptoe reacted to the culture filtrates by forming necrotic lesions with dark margins and a dark necrotic center, which resembled lesions that are typically induced by the necrotrophic pathogen *B. sorokiniana* that contain phenolics accumulation. To determine if the *nec3* phenotype was possibly induced by a pathogen derived proteinaceous effector, pronase treatment of the culture filtrates were performed prior to leaf infiltrations. The pronase treated culture filtrates resulted in no observable phenotypic change compared to the culture filtrates without pronase treatment (S1 Figure).

### DAB Staining and Electron Microscopy

The *nec3* mutants develop lesions that expand more rapidly and were larger than those of the WT genotypes post *B. sorokiniana* infection, thus, it was expected that the mutants would express differential ROS production. DAB staining associated with most of the *B. sorokiniana* penetration sites were observed as early as 12 hpi on *nec3-γ1* inoculated leaves (Figure 4). The DAB staining associated with pathogen penetration sites were only observed in the resistant WT Bowman line after 18 hpi. During the later time-points (24-36 hpi) the DAB staining associated with *B. sorokiniana* penetration and colonization in the susceptible *nec3-γ1* mutant (Bowman background), showed rapid spread of ROS to neighboring host epidermal cells, as well as underlying mesophyll cells. However, in the resistant WT Bowman seedlings, the DAB staining that started to appear at ~ 24 hpi had a higher intensity but remained limited to a few cells adjacent to the penetration site and did not expand at the later time-points during the infection process as was observed with the *nec3-γ1* mutant (Figure 4)

**Figure 4.**
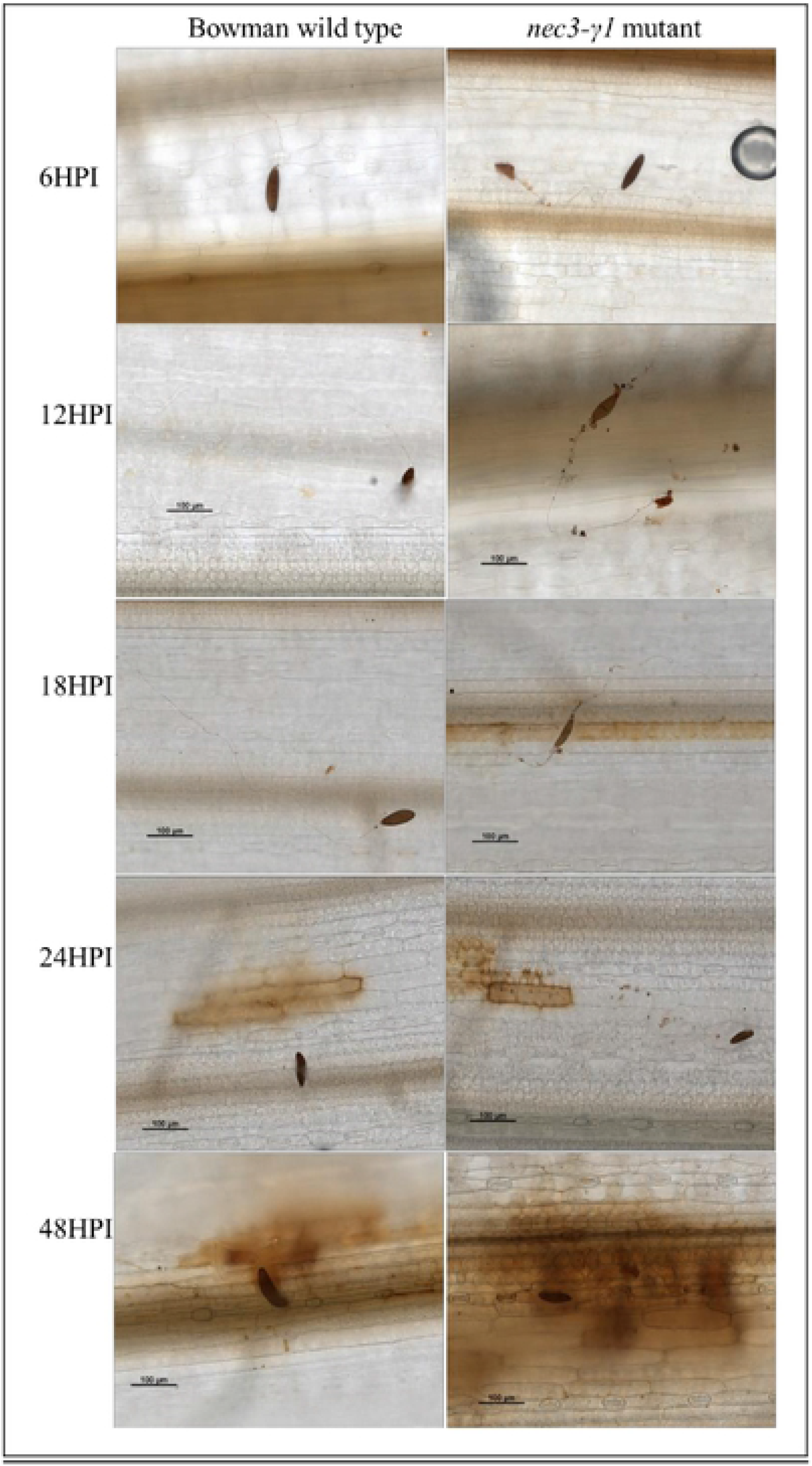
Microscopic visualization for comparison of the barley *nec3-γ1* mutant and Bowman wildtype for ROS production and pathogen growth on the leaf surface during infection process. On the left is the Bowman wildtype and on the right is the *nec3-γ1* mutant ROS production during infection of *Bipolaris sorokiniana* isolate ND85F at 6, 12, 18, 24, 48 hours post inoculation (HPI) where multiple branching of germ tubes, localized HR, and mutant specific interaction with the cuticle/of the barley leaves is observed as early as 12 hours post inoculation.

Time course light microscopy following DAB staining post *B. sorokiniana* isolate ND85F inoculations showed normal spore germination and germ tube growth on WT Bowman with germ tubes growing from each end of the conidium and little to no branching (Figure 4 and 5). Spore germination on the *nec3-γ1* mutant appeared normal, however aberrant germ tube branching was consistently observed compared to the growth on WT Bowman (Figure 4 and 5). Also, after DAB staining of the *nec3-γ1* mutant it was also consistently observed that dark colored debris would accumulate along the path of the germ tube growth across the leaf surface (Figure 4B) compared to the growth on WT Bowman (Figure 4A). As it could not be discerned if the debris accumulating along the germ tube growth on the *nec3-γ1* mutant was host or pathogen derived, electron microscopy was used to generate higher resolution images during the early infection process. These images showed that the debris was derived from the host cuticle peeling away from the leaf surface around the region of germ tube contact with the leaf surface (Figure 5).

**Figure 5.**
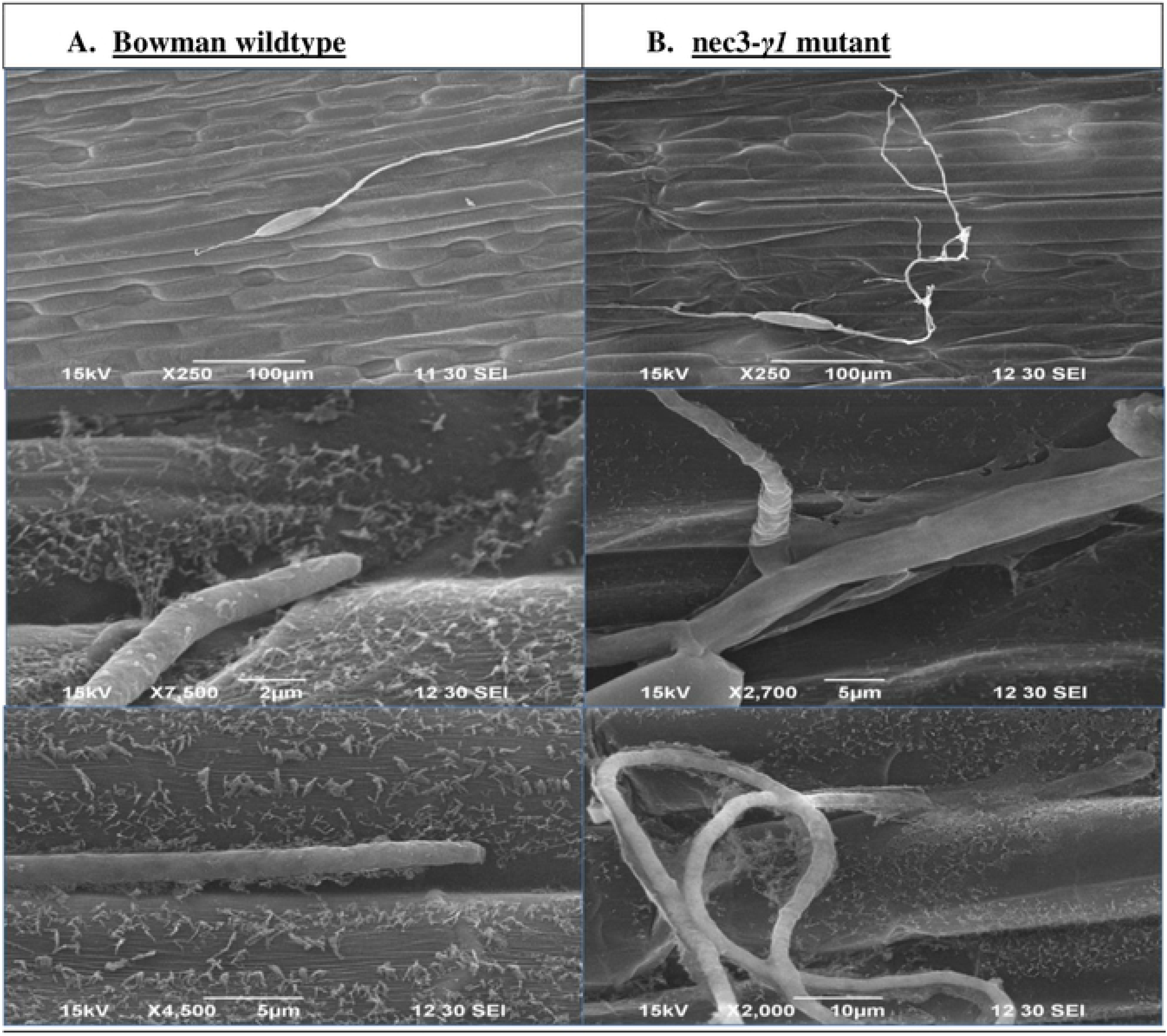
Electron Micrograph of the pathogen *Bipolaris sorokiniana* growth on the leaf surface of the barley *nec3-γ1* mutant and Bowman wildtype during the infection process.5A. On the left is the Bowman wildtype with normal *B. sorokiniana* isolate ND85F growth and intact cutin layer of host at 12 hours post inoculation. 5B. On the right is abnormal growth of *B. sorokiniana* isolate ND85F with multiple branching of germ tubes and mutant specific interaction with the cuticle of the barley leaves at 12 hours post inoculation.

### nec3 Genetic Map Development

A genetic map of the *nec3* gene was developed using homozygous F_2_ recombinant progeny from the cross between *nec3-γ1* and Quest (*γ1* x Q). Homozygous mutant individuals were identified by inducing the *nec3* phenotype with *B. sorokiniana* isolate ND85F. Due to the typical recessive nature of mutant genes, a 3:1 wild type: mutant ratio was expected. Unfortunately, a lethal chlorophyll-/albino background mutation killed 54 of the 200 F_2_ progeny assayed. The albino mutation was not linked to *nec3* and segregated in a recessive 3:1 single gene manner (χ^2^ = 0.32). After accounting for the albino plants that died 33 of the 146 surviving plants developed the *nec3* phenotype fitting the expected 3:1 ratio (χ^2^=0.17), indicating that *nec3* was segregating as a single mutant gene in a recessive manner.

To map the *Nec3* gene, the chromosome 6H region was saturated with molecular markers. According to the IPK cv Morex genomic sequences and POPSEQ positions (57,58), the *Nec3* gene co-segregated with marker GBM1212 located at 49.07cM on chromosome 6H. The proximal flanking marker GBM1423 was positioned at 49.22cM, 0.15cM proximal of GBM1212 and the distal flanking marker GBM1053 was positioned at 53.47cM, 4.4cM distal of GBM1212 (Figure 3) (58). Thus, the *Nec3* gene was delimited to a relatively large genetic interval of 4.55 cM. To further delimit the *Nec3* region an additional 29 SNP markers were genetically anchored to the *nec3* region of barley chromosome 6H using the *γ1* x Q population. The positions of the 29 SNP markers on the genetic map were in perfect linear order with the WGA Morex sequence released in 2019 (Monat et al., 2019) and the barley POPSEQ positions. The physical *nec3* region was flanked by the SNP marker SCRI_RS_171247 distally at pseudomolecule position 39.5 Mb and marker SCRI_RS_239962 proximally at 56.0 Mb on chromosome 6H, delimiting the region to ~ 0.14 cM based on the barley POPSEQ positions. This genetic interval correlated to a physical region spanning ~ 16.5 Mb containing 149 high confidence annotated genes according to the newly release barley genome sequence and annotation (Figure 6).

**Figure 6.**
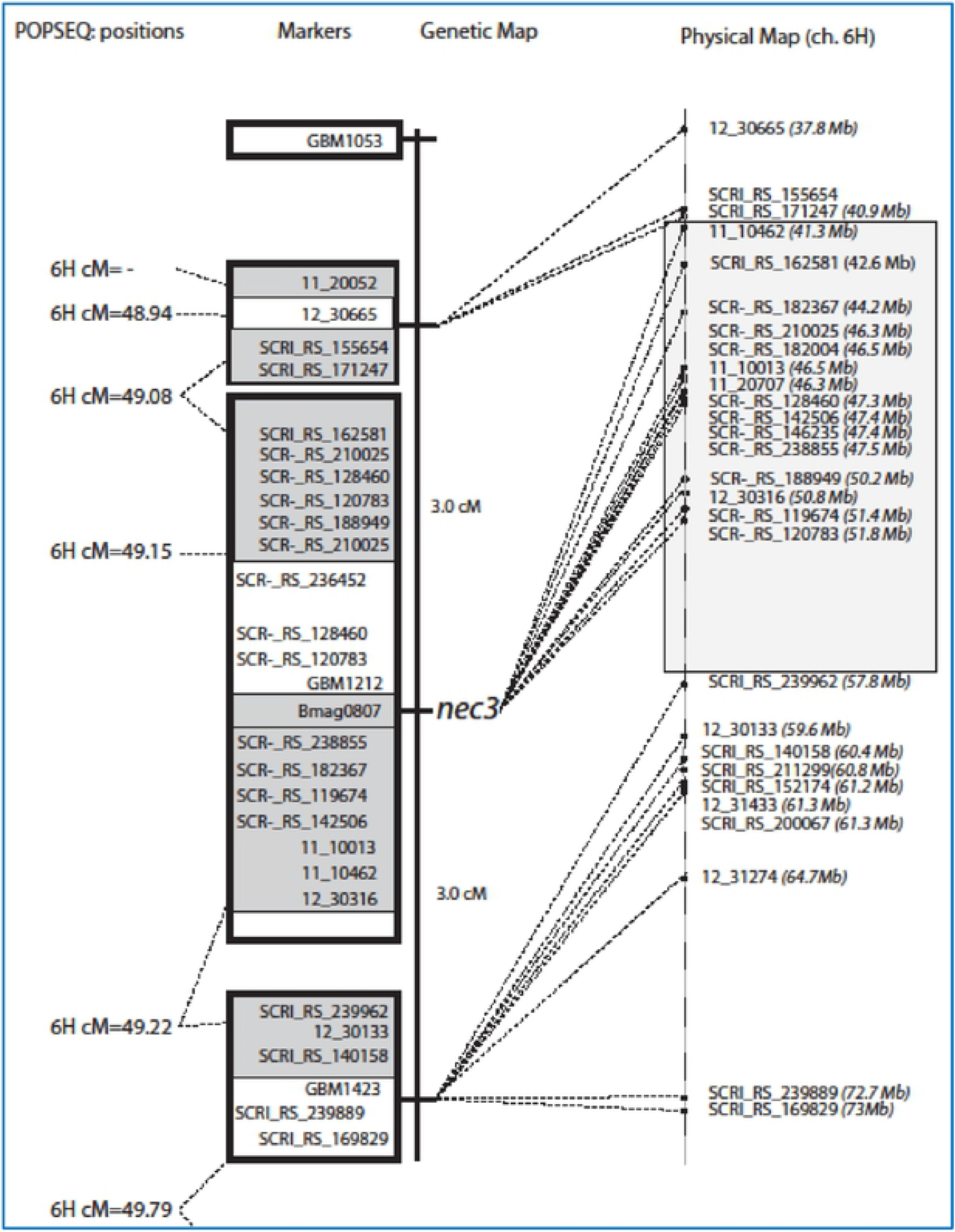
Genetic and physical map of the *nec3* region. On the left is the genetic map developed from 33 homozygous mutant F_2_ individuals (representing 66 recombinant gametes) from the cross between *nec3-γ1* and Quest. The genetic distances, based on recombination frequency is shown with the boxes indicating cosegregating markers based on the F_2_ map and white or gray shading indicating cosegregating markers based on POPSEQ consensus positions, which are given on the far left. The relative physical location of the markers is shown on the right which were derived from the new released whole genome assembly from the IPK database.

### Exome Capture for Identification of Candidate Genes

Sequences obtained from WT Bowman, WT Steptoe, *nec3-γ1* (Bowman background), *nec3.l* (Steptoe background), and *nec3.m* (Steptoe background) mutants generated after exome capture using the barley exome capture array were analyzed to identify potential deletions within the 149 high confidence candidate genes in the *nec3* region (58). A total of 11 genes in the region were not represented in the exome capture probe set, as shown in S3 Table. The 138 annotated high confidence genes (S2 Table) present in the exome probe set and captured in the *nec3* region were analyzed and no deletions were identified from the three independent mutants. To address the pitfall of the 11 uncharacterized gene missing in the exome capture, RNAseq analysis was carried out to analyze genes that were missed by the exome capture analysis.

### RNAseq and Candidate Gene Identification

An RNAseq analysis was performed on the *nec3-γ1* mutant and WT Bowman post inoculation with *B. sorokiniana* isolate ND85F to analyse the 11 genes in the delimited region missing from the exome capture analysis. The RNAseq also allowed for a thorough analysis of the global regulation of the transcriptomes in WT Bowman and the *nec3-γ1* mutant generated in the Bowman background upon *B. sorokiniana* isolate ND85F inoculation. In order to identify candidate genes, the total reads obtained and percentage alignment to the Morex reference genome were analysed along with comparative analysis between mutant and WT Bowman reads (S4Table) (GenBank Bio project PRJNA666939). The RNAseq analysis at 72 hpi identified a total of 10,473 differentially expressed genes (DEGs) with greater than a threefold change (5,171 upregulated and 5,303 downregulated) in the *nec3-γ1* mutant compared to the non-inoculated *nec3-γ1* mutant (Accession SRR12763054-SRR12763062). In resistant WT Bowman, 5,463 DEGs (2,803 upregulated and 2,661 downregulated) were identified in comparison to the non-inoculated WT Bowman control. Interestingly, the comparison between Bowman and *nec3-γ1* mutant transcriptome profiles during the *B. sorokiniana* isolate ND85F infection process revealed 3 genes upregulated in Bowman or downregulated in *nec3-γ1* mutant and nine genes that were downregulated in Bowman or upregulated in *nec3-γ1* mutant as shown as a venn-diagram and table (S4 Figure, S4 Table). The comparative analysis of the RNAseq data for the 11 genes missing from the exome capture probe set identified a 13 nucleotide deletion in the predicted exon1 of the cytochrome P450 gene HORVU.MOREX.r2.6HG0460850 in the *nec3-γ1* mutant (Figure 7). Eight of the eleven genes missing from the exome capture probe were not differentially expressed at control or in the pathogen induced samples at the tested time-point in both WT Bowman or the *nec3-γ1* mutant (S3 Table, fold changes reported as non-significant (NS)). Interestingly, the candidate *Nec3* gene was upregulated 1,126 fold in susceptible *nec3-γ1* mutant after pathogen inoculation and only upregulated 118 folds in WT Bowman after pathogen inoculation (S3 Table).

**Figure 7.**
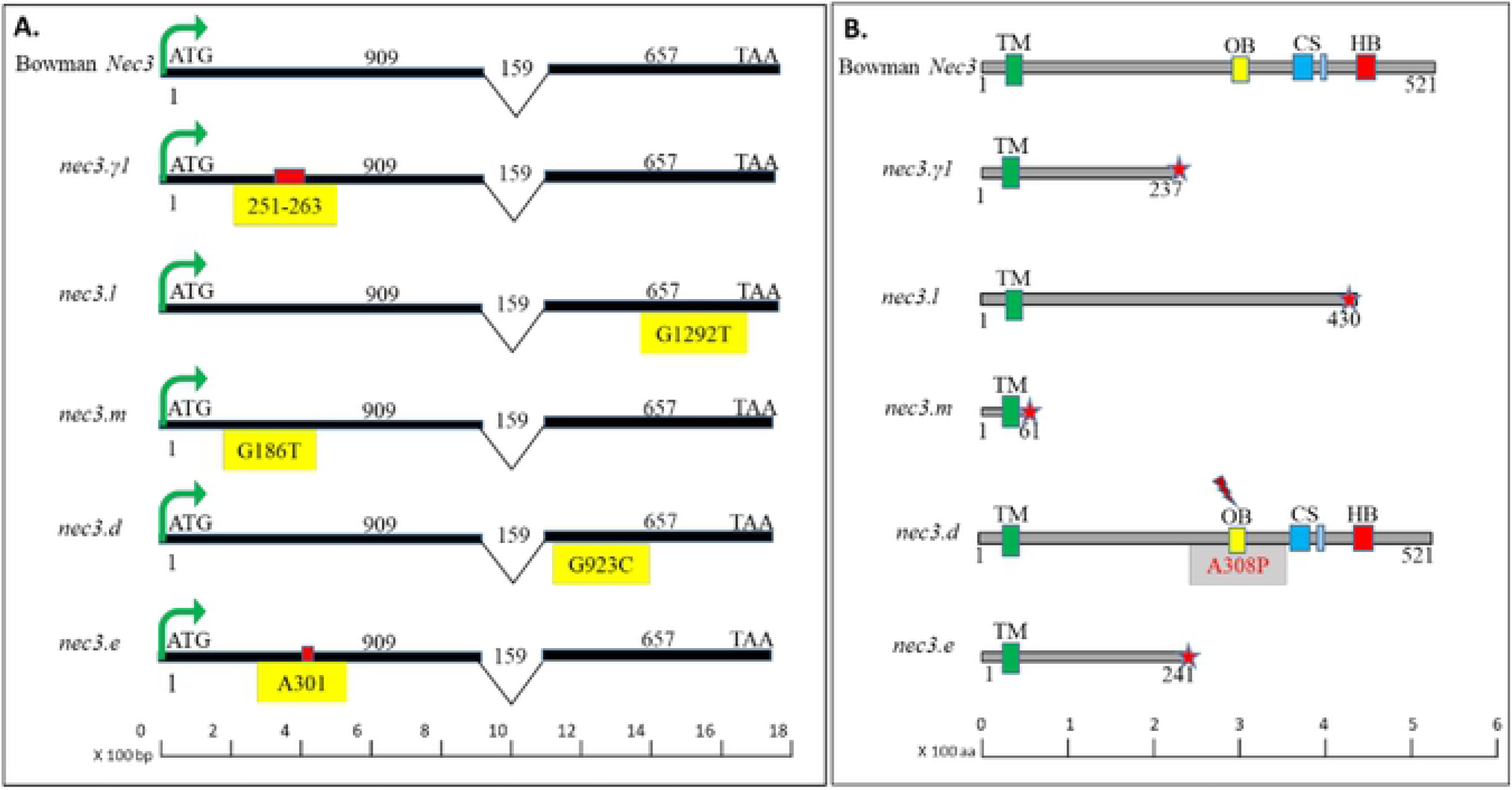
The allele analysis and protein polymorphism of the barley *nec3* mutants and Bowman wildtype. 8A. The genomic and cDNA structures for the barley *Nec3* gene in bowman wildtype and *nec3* mutants are shown from top to bottom (Bowman wildtype, *nec3-γ1*, *nec3*.l, *nec3*.m, *nec3*.d and *nec3*.e), where genomic and predicted mRNA structures are represented to scale with exon (black), intron (black Vs), start codon (ATG) and stop codon (TAA) and the mutations are denoted in the yellow box, where deletions are represented by red box above them. 8B. The barley *Nec3* protein structure in bowman wildtype and *nec3* mutants are shown from top to bottom (Bowman wildtype, *nec3-γ1*, *nec3*.l, *nec3*.m, *nec3*.d and *nec3*.e), where gray represent protein length, green bar represents transmembrane domain (TM), yellow bar represents oxygen binding (OB) and activation conserved residue AGxDTT, the blue bars represent the clade signatures (CS) with conserved residues ExxR and P(E)R(F) and red bars represent the heme binding (HB) with conserved residues as FxxGxRxCxG of the p450 clade. The star shows the truncated protein in the *nec3-γ1*, *nec3*.l, *nec3*.m, and *nec3*.e mutant and *nec3.d* has a A308P substitution represented by red lightening sign in the conserved residue of Oxygen binding domain of AGxDTT to PGxDTT, in the logos of the p450 protein family in plants.

### Cytochrome P450 Mutant Allele Analysis

The candidate *Nec3* Cytochrome P450 gene (HORVU.MOREX.r2.6HG0460850) was the only gene identified in the *nec3* region that contained a mutation in *nec3-γ1.* Analysis of the other four independent *nec3* mutants via PCR amplicon sequencing compared to their respective WT genotypes showed that all five alleles contained mutations that would disrupt the function of the predicted translated protein. The HORVU.MOREX.r2.6HG0460850 gene encodes a 1,566 bp gene (supported by RNAseq data) predicted to be translated into a 521 aa protein with an N-terminal transmembrane domain and a C-terminal p450 domain. The p450 domain has three conserved motifs, the oxygen binding (AGxDTT), clade signatures (ExxR) (P(E)R(F)) and heme binding (FxxGxRxCxG) motifs (Figure 7) (59). The *nec3-γ1* mutant has a 13-nucleotide deletion at position 251 to 263 bp in exon1, that is predicted to result in a premature stop codon and truncated protein of 237 aa, eliminating the entire p450 domain (Figure 7). A G186T nucleotide conversion in *nec3.m* was predicted to produce a premature STOP codon and a truncated protein of 61 aa eliminating the p450 domain (Figure 7). The *nec3.e* mutant contained the single nucleotide deletion A301 that was predicted to cause a frame shift resulting in a premature STOP codon and predicted truncated protein of 241 aa also eliminating the p450 domain (Figure 7). The G1292T nucleotide conversion in the *nec3.l* mutant was predicted to produce a premature STOP codon at aa position 430, eliminating the heme-binding motif present in the p450 domain (Figure 7). Intriguingly, the allelic *nec3.d* mutant that produces an atypical *nec3* phenotype that contain dark phenolics and a relatively smaller lesion size compared with the other *nec3* mutants had a G923C nucleotide substitution mutation, which resulted in the predicted A308P aa conversion present in the conserved residues of the oxygen binding (AGxDTT) motif of the p450 domain, yet is still predicted to produce a full length possibly partially functional protein (Figure 7).

### Exogenous Supplementation of DAMPs, Serotonin and Tryptamine

Infiltrations of WT Bowman, *nec3-γ1*, WT Steptoe and *nec3.l* with the known DAMP trigalacturonic acid was carried out at three concentrations (10mg/mL, 1mg/mL and 0.1mg/mL), the bacterial PAMP FLG22 at a concentration of 1mg/mL, and chitin (100μg/ml) resulted in no observable reactions up to one week after infiltration (S2 Figure), compared to the mock controls treated with water. To test if the *nec3* mutant gene is also deficient in serotonin biosynthesis we tested if exogenous root feeding of serotonin would produce changes in the size and shape of necrotic lesions similar to what was observed for the *sl* mutant in rice. It was observed that the serotonin fed *nec3-γ1, nec3.d, nec3.e*, *nec3.l* and *nec3.m* mutants and wild type plants displayed significantly smaller lesion and had little to no observed chlorosis surrounding the lesions on the secondary leaves at seven days post pathogen inoculations compared to the mock water controls (S3 Figure A and B). However, the tryptamine fed plants had increased lesion sizes for *nec*.d and the WT seedlings compared to the water controls with significantly larger lesions for all genotypes compared to the serotonin fed plants (S3 Figure A and B).

## DISCUSSION

The cytochrome P450 gene (HORVU.MOREX.r2.6HG0460850) was identified as the *Nec3* gene. Barley *nec3* mutants that produce the atypical and distinct *nec3* phenotype, large cream to orange necrotic lesions lacking the dark pigmentation indicative of the accumulation of serotonin phenolic compounds, were characterized in the cv Bowman, Steptoe, Proctor and Villa backgrounds. Four of the independent mutants had deletions or nucleotide substitutions in the *Nec3* cytochrome P450 gene that resulted in predicted nonfunctional truncated proteins. Interestingly, a fifth independent mutant shown to be allelic to *nec3* produced the typical DLMM phenotype with smaller dark necrotic lesions containing serotonin or phenolics accumulation (Figure 1). This *nec3* mutant with a single predicted A308P aa conversion in the conserved oxygen binding motif of the *Nec3* p450 domain represents an excellent functional tool for future investigation of the role that serotonin (31) or phenolics accumulation in necrotic lesions play in sequestration of lesion expansion or pathogen colonization during HR responses. It was also discovered that *nec3* is not a true DLMM as it is only expressed when the barley mutants are challenged by diverse specialized pathogens.

Originally the *nec3* mutants were classified as lesion mimic mutants (LMMs) (52), referred to as disease lesion mimic mutants (DLMMs) here, that spontaneously develop necrotic lesions when they reach a certain developmental stage. However, contrary to these observations, the *nec3* phenotype is only expressed when elicited by a diverse taxonomy of pathogens. The *nec3* phenotype is possibly triggered through PAMP elicitor recognition such as chitin subunits released from fungal pathogens during pathogen entry and host colonization. However, our experiments determined that the known DAMP trigalacturonic acid the bacterial PAMP FLG22 and the fungal PAMP chitin didn’t induce the *nec3* phenotype. Previous research and observations that led to the description of *nec3* as a DLMM were conducted under less controlled greenhouse and field environments, where the *nec3* phenotype was consistently expressed without apparent biotic or abiotic stress induction (54). However, in the same study Kesia et al. (2010) observed that mutant seedlings grown in growth chambers for transcriptome analysis did not exhibit the *nec3* phenotype. In our screen of a variety Bowman mutant population for mutants of the dominant *B. sorokiniana* isolate ND90Pr susceptibility gene (56), the *nec3-γ1* mutant was identified. The *nec3-γ1* mutant exhibited the phenotype consistent with the previously described and characterized *nec3* mutants (60), however, the phenotype appeared after inoculation with the pathogen leading us to hypothesize that the *nec3* characteristic lesions were induced by pathogen challenge. To test this hypothesis a sterile environmental condition experiment was conducted that determined the *nec3* phenotype was induced in all the *nec3* mutants (*nec3-γ1, nec3.d, nec3.e nec3.l*, and *nec3.m*,) by the necrotrophic ascomycete fungal pathogens *B. sorokiniana*, *P. teres* f. *teres*, and *P. teres* f. *maculata*, as well as the biotrophic ascomycete *Blumeria graminis* (Figure 2). During host penetration, these fungal pathogens are known to form appressoria-like structures and penetration pegs that puncture the cell wall and plasma membrane causing cellular damage (61–63). However, a cell damage assay using mechanical disruption of barley leaves also did not induce the *nec3* phenotype. Host cell disruption and pathogen penetrative structures are also accompanied by proteinaceous and secondary metabolite effector molecules that induce host immune responses that induce temporally regulated and spatially confined PCD responses.

Interestingly, the *nec3* phenotype was not elicited by either virulent or avirulent races of the basidiomycete biotrophic pathogen *P. graminis* f. sp. *tritici*, whose strategy is “incognito” host entry through the stomata. It is hypothesized that *P. graminis* f. sp. *tritici* hijacks the stomatal aperture regulation of the cereal hosts in order to enter through the natural openings at night without being detected (64,65). The *nec3-γ1* mutant in Bowman background contains the stem rust resistance gene *Rpg1* (66,67), which elicits an effective race specific immunity response yet did not elicit the *nec3* phenotype. Recent research has reported that *Rpg1*-mediated defenses are non-HR responses (64,68), thus suggesting that the *nec3* phenotype may still be elicited by HR-mediated resistance responses.

The bacterial pathogen *X. translucens*, which causes tissue damage by employing the penetrative Type-III secretion system to deliver virulence effectors also induced the *nec3* phenotype. Bacterial pathogens also utilize enzymes and other molecules to increase the permeability of the cell wall. This leakiness allows nutrients to be acquired by the bacteria facilitating colonization. Collectively, it is possible that the loss of cellular integrity induced by the bacterial pathogen induces the *nec3* phenotype or a PAMP-like bacterial elicitor also triggers the *nec3* phenotype. However, our experiments also determined that the bacterial PAMP flg22 didn’t induce the *nec3* phenotype. Thus, it appears that fungal or bacterial PAMPs as well as mechanical damage are not sufficient to induce the *nec3* phenotype suggesting a more complex host-pathogen interaction is required.

Necrotrophic fungal pathogens produce proteinaceous, non-proteinaceous, and secondary metabolite effectors that initiate host PCD in order to colonize the dead and dying tissue via necrotrophic effector triggered susceptibility (NETS) (24,25,69,70). To determine the nature of the elicitor/s of the *nec3* response a pathogen culture filtrate infiltration assay was performed. *B. sorokiniana* crude culture filtrates elicited the distinct *nec3* PCD phenotype on the mutants and characteristic dark pigmented lesion in the susceptible barley line Steptoe, but not in the resistant cv Bowman (S1 Figure). However, in the *nec3.d* mutant, culture filtrates didn’t produce the pronounced dark pigmented lesions that were observed with the *B. sorokiniana* isolate ND85F inoculations. These data suggested that the differential phenotype on this mutant during pathogen colonization is dependent on effectors or interaction occurring during pathogen colonization that are not present in the pathogen culture filtrates. Pronase treatment of the culture-filtrate did not abolish the elicitation of the distinct *nec3* lesions at the infiltration site. Thus, we speculate that the pathogen effector/s in the culture filtrates that elicit the *nec3* phenotype is/are not proteinaceous effector/s but rather represent a PAMP such as chitin, a secondary metabolite, a non-proteinaceous toxin, an RNA molecule or a tightly folded small protein. It cannot be ruled out that *nec3* is involved in suppressing PCD responses elicited by DAMPs including cell wall components, eATP, eDNA or other endogenous molecules disrupted during the infection process as recognition of cellular damage elicits defense responses across all multicellular life including algae, fungi, fish, insects, mammals and plants (71). However, our experiments showed that the oligogalacturonide residue (OG) extracellular matrix subunits trigalacturonic acid did not elicit the *nec3* phenotype.

To identify *nec3* a genetic map was generated with F_2_ recombinant progeny from the cross between *nec3-γ1* (Bowman background) and the barley variety Quest (referred to as the *γ1* x Q population). The *nec3* region was delimited to ~0.14 cM on barley chromosome 6H between the flanking markers SCRI_RS_155654 and SCRI_RS_239962. The genetic mapping delimited *Nec3* to a physical distance of ~16.49 Mb on barley chromosome 6H containing 149 high confidence (HC) annotated genes based on the recently refined assembly and annotation of the barley Morex reference genome (2019). Exome capture sequencing of WT Bowman, WT Steptoe and three *nec3* mutants (*nec3-γ1, nec3-l* and *nec3-m*) was generated for comparative analysis. However, the exome capture library only contained probes for 138 of the 149 annotated HC genes present in the *nec3* region. The exome capture analysis did not identify any mutated genes within the *nec3* region but the identification of 11 HC genes in the region that were missing from the exome capture probe suggested the need for further comparative analysis using an RNA sequencing approach. The RNAseq comparative analysis identified a 13 bp deletion in the HORVU.MOREX.r2.6HG0460850 HC gene model predicted to encode a Cytochrome P450 gene in the *nec3-γ1* mutant. Utilizing PCR amplification and Sanger sequencing mutations were identified in all four independent mutants that produce the typical *nec3* phenotype (*nec3-γ1*, *nec3*.e, *nec3*.*l*, and *nec3*.*m*) that were predicted to result in nonfunctional truncated proteins. The HORVU.MOREX.r2.6HG0460850 allele in the fifth *nec3* mutant, *nec3.d*, that expresses the atypical dark necrotic spots (Figures 1 and 8) contained a critical amino acid substitution in the *nec3.d* allelic mutant.

The genetic mapping and presence of irradiation induced deletions and nucleic acid substitutions in five independent mutants in four different genetic backgrounds confirmed that HORVU.MOREX.r2.6HG0460850 is the functional *Nec3* gene. The *Nec3* gene model was predicted to encode a 521 amino acid Cytochrome P450 family protein. Cytochrome P450s are heme-thiolate proteins representing one of the largest super families of proteins with enzymatic activity. Within the superfamily the amino acid sequence conservation is low and only three residues belonging to the conserved sequence motif (CSM) are absolutely conserved. However, the general topography and structural folding are conserved across the members (72). Only the CYP51 gene family of P450s is conserved among plant, fungi and animal phyla with the presence of CYP51 orthologs in the bacterium *Mycobacterium tuberculosis*, possibly due to lateral gene transfer (59). Sequence comparison and phylogenetic analyses of the NEC3 protein among the monocots placed it in the CYP71 clan as a cytochrome P450 71A1-like proteins (http://www.p450.kvl.dk/blast.html) suggesting involvement in plant lineage specific metabolism (73) (S5 Figure).

Interestingly, the Sekiguchi lesion (*sl*) mutant in rice produces orange tan necrotic lesions, similar to the barley *nec3* phenotype. The *sl* gene was identified via map-based cloning and shown to encode CYP71P1 in the cytochrome P450 monooxygenase family that shares 90.1 % amino acid similarity and 86.7% identity with the barley *Nec3* protein (S6 Figure) (31). The *sl* gene encodes a Tryptamine 5-Hydroxylase (T5H) enzyme that converts tryptamine to serotonin in plants in the shikimate pathway (31). The rice *sl* gene mutants were susceptible to the rice blast fungus (*Magnaporthe grisea*) and rice brown spot fungus (*Bipolaris oryzae*) and the *sl* necrotic lesions were also induced by the N-acetylchitooligosaccharide (chitin) elicitor (55). The *sl* mutant susceptibility to rice blast was eliminated by exogenous application of serotonin in rice leaves (55) and the exogenously applied serotonin was deposited into the cell wall of the *sl* lesion tissue restoring the dark pigmentation and resistance to the necrotrophic pathogen *B. oryzae* (31).

Annotation of the NEC3 protein sequences from the five allelic *nec3* mutants showed that the four with the typical *nec3* phenotypes encoded premature stop codons due to the induced mutations. The *nec3.d* mutant with the smaller dark pigmented lesions contained a single nucleotide substitution that resulted in an A308P amino acid conversion in the predicted oxygen binding and activation domain of the conserved P450 residues (Figure 7). The data showed that upon pathogen infection, the *nec3.d* allele was induced and predicted to be translated into a full-length protein with a nonfunctional oxygen binding domain due to the single amino acid substitution (Figure 7). Interestingly, the *nec3.d* mutant although allelic to the truncated nonfunctional *nec3* alleles expresses comparatively smaller dark pigmented lesions presumed to be due to higher serotonin/phenolics accumulation. We attributed the smaller lesion size of *nec3.d* compared to the typical cream colored *nec3* phenotypes expressed by *nec3-γ1*, *nec3*.e, *nec3*.*l*, and *nec3*.*m* mutants to be due to the A308P amino acid conversion, resulting in altered P450 function. Thus, the *nec3.d* mutant containing the partially functional full-length protein may be able to convert Tryptamine to Serotonin but due to the critical A308P substitution has a leaky control on PCD and is not able to keep it in check upon pathogen recognition. Evolutionarily, plants adapted to utilize oxidative polymerization of serotonin for the modification of cell walls as part of the physical barrier and phenolics deposition at the infection site during pathogen infection to inhibit pathogen growth and sequester them in the foci of dead cells. This mechanism may also act as a signaling mechanism to sequester lesion expansion to preserve the leaf’s photosynthetic capability after activating PCD-mediated defenses in response to pathogen challenge.

To rescue the serotonin deficiency and determine the effects of excess tryptamine, the serotonin precursor, barley *nec3* mutants and wildtype seedlings were provided with exogenous serotonin and tryptamine via root feeding. The exogenous application of serotonin reduced the pathogen induced lesions in the *nec3* mutants rescuing the phenotype induced by the apparent deficiency caused by the nonfunctional *Nec3* genes. Interestingly, the exogenous feeding with tryptamine increased the lesion sizes in wildtype and nec3.d mutant implicating NEC3 in the conversion of tryptamine into serotonin. Thus, we hypothesize that *Nec3* also encodes a tryptamine 5-hydroxylase similar to the *sl* gene in rice and exogenous application of upstream or downstream substrates in the pathway modulate the final phenotypic outcomes. These observations and apparent phenotypic differences described above suggest that the oxygen binding domain plays a critical role in the function of NEC3 regulation of PCD as well as the pathway leading to serotonin (31) accumulation. The non-functional *nec3* mutant alleles predicted to produce truncated proteins failed to regulate the PCD response, resulting in expanded necrotic lesions with no defined border. However, the *nec3.d* mutant still accumulates phenolics and has a defined lesion border that sequesters the expansion of the lesion in the presence of the pathogen. This phenomenon is also observed in resistant WT Bowman.

The RNAseq analysis showed that the barley *NEC3* gene is highly upregulated in WT Bowman and the *nec3-γ1* mutant, 72 hours post *B. sorokiniana* inoculation, thus the upregulation of the *sl* ortholog suggests an increased need to convert Tryptamine to Serotonin in barley because of its importance in regulating PCD-mediated immunity. The question of how or why serotonin/phenolics accumulate in necrotic lesions still needs to be answered and the *nec3.d* allele presents a unique tool for functional characterization of the role of serotonin/phenolic deposition in pathogen sequestration or the regulation of lesion expansion.

Necrotrophic fungal pathogens produce proteinaceous, non-proteinaceous, and secondary metabolite effectors that initiate host PCD in order to colonize the dead and dying tissue via necrotrophic effector triggered susceptibility (NETS) (24,25,69,70). To determine the nature of the elicitor/s of the *nec3* response a pathogen culture filtrate infiltration assay was performed. *B. sorokiniana* crude culture filtrates elicited the distinct *nec3* PCD phenotype on the mutants and characteristic dark pigmented lesion in the susceptible barley line Steptoe, but not in the resistant cv Bowman. However, in the *nec3.d* mutant, culture filtrates did not produce pronounced dark pigmented lesions as seen with the *B. sorokiniana* ND85F isolate inoculation on *nec3.d*. This suggests that this single amino acid substitution allows for the induction of enhanced PCD elicitation by the pathogen in the WT Bowman background yet possibly still retains partial function allowing for serotonin metabolism and build up in the lesions which still functions to sequester the pathogen and necrotic lesion expansion. Interestingly, the culture filtrate infiltration of the *nec3.d* mutant was indistinguishable from the other *nec3* mutants with no dark pigmentation in the absence of the pathogen. This suggested that post-elicitation, the *nec3.d* mutant behaves similarly to the other non-functional mutants but in the presence of the pathogen it is still able to mount some level of serotonin biosynthesis and suppression of lesion expansion and pathogen growth.

The pronase treatment of the culture-filtrate did not abolish the elicitation of the distinct *nec3* lesions at the infiltration site. Thus, we can speculate that the pathogen effector/s in the *B. sorokiniana* culture filtrates that elicit the *nec3* phenotype is not a proteinaceous effector but similar to the *sl* mutant in rice may be elicited by a PAMP, possibly chitin as it elicited the *sl* phenotype. However, none of the tested known PAMPs; bacterial FLG22 or fungal chitin or the DAMP triglyciraldehide elicited the *nec3* response.

In Arabidopsis the CYP84A1 EMS mutant was shown to have altered lignin composition (74). Thus, alterations of the cell wall composition could affect the pathogen’s spatiotemporal interactions with the host at the leaf surface which we tested microscopically in the *nec3-γ1* mutant. Upon microscopic observations of *B. sorokiniana* growth pattern on the *nec3* mutant compared to WT Bowman it was observed that the pathogen’s infection hyphae interaction with the host cuticle was abnormal showing that the cuticle was unstable and peeled away from the *nec3-γ1* leaf surface where there was contact (Figure 5B). This aberrant interaction apparently disrupted the signaling in the pathogen’s infection hyphae. When growing across the *nec3* mutant leaf surface the *B. sorokiniana* hyphae branched profusely and produced surface debris apparently due to unstable cuticle along the germ tubes (Figure 5). The profuse branching of pathogen and plant cell surface destabilization could result in many host-pathogen contact points with the *nec3* mutants allowing the plants to recognize more pathogen effector molecules. This aberrant interaction possibly lead to the rapid ROS production in the *nec3* mutants and the runaway cell death due to the lack of serotonin accumulation which may play a role in the sequestration of PCD mediated lesion expansion similar to the *sl* mutant in rice. The DAB assays supported this conclusion as the *nec3-γ1* mutant produced ROS when infected by the necrotrophic pathogen *B. sorokiniana* as early as 12 hpi as compared to a delayed ROS in the susceptible Steptoe and resistant Bowman which was detected at 18 and 24 hpi, respectively (19) (Figure 4).

Here we report on the identification of the barley *Nec3* gene as a P450 CYP71A1 cytochrome oxidase that plays a role in suppressing PCD responses that are initiated by diverse adapted barley pathogens. We hypothesize that *Nec3* is a negative regulator of the PCD and is an ortholog of the *sl* gene identified and characterized in rice which plays a role in serotonin biosynthesis and buildup in necrotic lesions. The barley *Nec3* gene provides a valuable resource to study the control of PCD and HR responses and our collection of mutant alleles especially the *nec3.d* mutant will be an excellent tool for determining the role of serotonin and possibly phenolic compound deposition in necrotic lesions for the sequestration of pathogen colonization or lesion expansion.

## MATERIALS AND METHODS

### Plant Material and Bipolaris sorokiniana Isolates

Approximately 0.25 kg of cv Bowman seed was irradiated using 20 kRADs of γ-radiation at the Washington State University nuclear reactor in Pullman, WA USA. The cv Bowman (Reg. no. 197) PI483237 was jointly developed by North Dakota State University, Fargo, ND and the USDA-ARS and released in 1984 (Fanckowiak et al., 1985). The M_1_ plants were allowed to self and M_2_ seed were bulked and ~ 8,000 M_2_ seedlings were screened by inoculation with *B. sorokiniana* isolate ND90Pr following the inoculation procedure described by Fetch and Steffenson (75). The *nec3-γ1* mutant described here was originally identified from this γ-irradiated M_2_ population when screening for mutants resistant to *B. sorokiniana* isolate ND90Pr to identify individuals with mutations in the putative cv Bowman dominant susceptibility gene (76).

The previously identified *nec3* mutants utilized in allelism tests and candidate gene allele analysis were irradiated using x-ray mutagenesis in the cv Proctor and Villa backgrounds and by fast neutron mutagenesis in the cv Steptoe background (54). The *nec3.d* (GSHO 1330) mutant was generated in cv Proctor (PI 280420) (52) and the *nec3.e* (GSHO 2423) mutant was generated in cv Villa (PI 399506) (51). The *nec3.d* and *nec3.e* mutants used in this study were near isogenic lines developed by recurrent backcrossing into the cv Bowman background. The *nec3.d* and *nec3.e* mutants were provided by Dr. Jerry Franckowiak and were *nec3.d* in Bowman*6 (BW629) and *nec3.e* in Bowman*6 (BW630). The *nec3.l* (GSHO 3605 formally known as FN362) and *nec3.m* (GSHO 3606 formally known as FN363) mutants were generated in the cv Steptoe (CIho 15229) background. The *nec3.1* and *nec3.m* mutant lines used in this study were provided by Dr. Andris Kleinhofs, Washington State University.

### Elicitation of nec3 Phenotype

An isolation box experiment was performed on the wildtype (WT) cv Bowman and *nec3-γ1* (Bowman background) plants over the course of a month to observe the development of the *nec3* phenotype independent of pathogen challenge. This experiment is thoroughly described in the supplementary information (S1 appendix).

### Pathogens that Induce nec3 Phenotypes

To determine which pathogens could induce the *nec3* phenotype, inoculations of the mutant line *nec3-γ1* and WT lines were performed using diverse fungal pathogens including *B. sorokiniana*, *Pyrenophora teres* f. *teres*, *Pyrenophora teres* f. *maculata*, *Pyrenophora tritici-repentis*, *Parastagonospora nodorum*, *Cercospora beticola*, and virulent and avirulent races of *Puccinia graminis* f. sp. *tritici*. The barley bacterial pathogen *Xanthomonas translucens* pv *undulosa* was also used to inoculate the plants. The method and materials followed in these experiments are thoroughly described in the S1 appendix.

### Infiltrations with DAMPs

Culture filtrate infiltrations for *B. sorokiniana* isolate ND85F were produced following the Liu et al., 2004 protocol and described in the S1 appendix (77). A syringe without a needle was used to infiltrate four secondary leaves of WT Bowman, WT Steptoe, *nec3-γ1*, WT Quest, *nec3*.*l* and *nec3*.*m*, *nec3*.d and *nec3*.e. The four treatments consisted of; 1) concentrated exudates + Fries Media, 2) concentrated exudates + MOPs buffer, 3) concentrated exudates + MOPs + Pronase (Sigma), and 4) Fries media + MOPS + Pronase. All treatments, concentrated exudates + Fries media, and concentrated exudates + MOPS were at 1:1 ratio with specifications performed according to Liu 2004 (78). Pronase treatments were performed at 1mg/mL and leaves were scored 4 and 7 days after infiltrations. Infiltrations were also performed with trigalacturonic acid, Flg22 and Chitin. Wildtype Steptoe and Bowman, *nec3.l* and *nec3-γ1* were infiltrated with the DAMP trigalacturonic acid at 10mg/mL, 1mg/mL and 0.1mg/mL, and with the PAMPs Flg22 at 1mg/mL and Chitin at 2μg/ml. All infiltrations with the controls were performed at the two-leaf stage. Following infiltration, plants were kept in a growth chamber with a 14-hour photoperiod at 22°C and 10 hours of dark at 18°C. Plants were observed every day up to seven days after infiltration.

### DAB Staining

To observe ROS at the site of infection, five 3 cm leaf samples were collected from secondary leaves at 6, 12, 18, 24 and 48 hpi from WT Bowman and *nec3-γ1* mutant seedlings inoculated with *B. sorokiniana* isolate ND85F. After detachment the leaves were immediately transferred to 10 ml of freshly prepared 1mg/ml DAB (Sigma Aldrich, MO) solution (pH 3.6) in 15 ml tubes following the protocol described by Solanki et al. (79).

### Electron Microscopy

Electron microscopy was performed on leaves collected from WT Bowman and *nec3-γ1* mutant seedlings at 12 hpi with *B. sorokiniana* isolate ND85F. The leaves were collected and cut into squares with a razor blade, fixed in 2.5% glutaraldehyde in sodium phosphate buffer (Tousimis, Rockville, Maryland USA) and stored at 4°C overnight. The sectioned leaf samples were rinsed with distilled water followed by rinsing with sodium phosphate buffer 1M solution at 7.4 pH and then dehydrated using eight washes of a graded alcohol series from 30% to 100% ethanol with incremental concentrations increased by 10% for each wash. The leaf samples were critical-point dried using an Autosamdri-810 critical point drier (Tousimis, Rockville, Maryland USA) with liquid carbon dioxide as the transitional fluid. The dried leaf samples were attached to aluminum mounts with silver paint (SPI Supplies, West Chester, Pennsylvania USA) and sputter coated with gold (Cressington sputter coater Redding, California USA). Images were obtained using a JEOL JSM-6490LV scanning electron microscope operating at an accelerating voltage of 15 kV.

### Allelism Crosses

The *nec3-γ1* mutant identified in this study was crossed with *nec3.1*, *nec3.m*, *nec3.d* and *nec3.e*. The resulting F_1_ seed were planted in the field with WT parental lines Steptoe and Bowman and phenotyped from seedling to adult plant stages. The F_2_ progeny were planted in the greenhouse and inoculated with the *B. sorokiniana* isolate ND85F and the *nec3* phenotype/disease was rated on the secondary leaves of the seedlings. The phenotyping was performed in the field and greenhouse to take advantage of the entire year. This was made possible due to the consistency of elicitation of the *nec3* phenotype when grown adjacent to susceptible spreader rows inoculated with the *B. sorokiniana* isolate ND85F in the field and when inoculated in the greenhouse with *B. sorokiniana* isolate ND85F.

### PCR-GBS Library Preparation, Ion Torrent Sequencing and SNP Calling

A PCR genotyping-by-sequencing (PCR-GBS) panel was developed using SNP source file sequences mined from the T3 database (80) as previously described (81). The POPSEQ positions were utilized from the IPK Barley BLAST Server (57) to identify 43 SNP markers that mapped to the *nec3* locus between the flanking SSR markers GBM1053 and GBM1423. The parental lines *nec3-γ1* (cv Bowman) and cv Quest plus the 33 F_2_ homozygous *nec3* mutant recombinants were assayed. The PCR-GBS library preparation, Ion Torrent sequencing and SNP calling were performed as described before (81,82). However, due to the low number of lines and markers the library was sequenced on an Ion Torrent PGM 314 chip. The genetic map in the delimited region was generated manually and the POPSEQ positions of each marker was mined from the IPK genome browser (https://webblast.ipk-gatersleben.de/barley_ibsc/viroblast.php).

### Nec3 Map Development

The original *nec3-γ1* M_2_ plant generated in the cv Bowman background was utilized as the female parent in a cross with the six-rowed cv Quest that was originally developed for malting and released by the University of Minnesota (83). Two hundred *nec3-γ1* x Quest F_2_ individuals were screened in the greenhouse for the *nec3* phenotype after inoculation with *B. sorokiniana* isolate ND85F as previously described and a chi square test was used to determine goodness of fit. Allelism tests were used to determined that *nec3-γ1* was a *nec3* mutant. To precisely map *nec3* 47 simple sequence repeat (SSR) and single nucleotide polymorphism (SNP) markers spanning the previously delimited *nec3* region near the centromere of barley chromosome 6HS were used to genotype the F_2_ individuals showing the homozygous *nec3* mutant phenotype. Unfortunately, a lethal chlorophyll/albino mutation killed 54 of the 200 F_2_ progeny assayed. This background mutation segregated in a recessive 3:1 single gene manner (χ^2^= 0.32) and based on segregation data was not linked to *nec3*, thus we were able to disregard the plants that died and use the remaining plants that developed the *nec3* phenotype to calculate the genetic ratio and map the *nec3* gene. Genomic DNA was extracted from *nec3-γ1*, Quest and the 33 homozygous *nec3-γ1* F_2_ progeny representing 66 recombinant gametes. The panel of 43 SNP markers (described above) and the four microsatellite markers (Bmag0807, GBM1053, GBM1212, and GBM1423) PCR amplified using oligonucleotide primers designed from the publicly available probe sequences mined from GrainGenes (84) were used to genotype the parental lines *nec3-γ1* (cv Bowman) and cv Quest and the 33 F_2_ recombinants (66 recombinant gamete). The data was used to construct the genetic map for the *nec3* region (S1 Table).

### Physical Map Development and Candidate Gene Identification

The 43 SNP markers were anchored to the barley genome using the IPK barley BLAST server (https://webblast.ipk-gatersleben.de/barley_ibsc/viroblast.php) with physical positions based on the 2019 Whole genome assembly of barley cv Morex. The genetic map anchored to the genome assembly was used to identify the physical region containing *Nec3*. The delimited region was used to identify high confidence annotated genes considered as candidate genes (58) (S2 Table).

### Exome Capture and Analysis

DNA was extracted from excised embryos of five germinated seeds of WT Bowman, WT Steptoe, *nec3-γ1* (Bowman background), *nec3.l* (Steptoe background), and *nec3.m* (Steptoe background) mutants. DNA extractions were performed on mechanically lysed samples using the PowerPlant Pro DNA isolation kit (Qiagen, CA), following the protocol described by Solanki et al. (82).

### RNAseq

The WT Bowman and *nec3*-*γ1* seedlings were grown for ~14 days until the secondary leaves were fully expanded in a growth chamber set at 14 hours light at 22°C and 10 hours dark at 19°C. The seedlings were inoculated with *B. sorokiniana* isolate ND85F following the procedure described earlier and the control seedlings were inoculated with water mixed with two drops of Tween20. Secondary leaf tissue was collected from three biological replicates (one individual seedling was considered as a biological replicate) from non-inoculated and inoculated seedlings at 72 hours post inoculation (hpi) and total RNA was extracted using the RNeasy Mini kit (Qiagen, CA). The RNAseq library was constructed using TruSeq RNA library prep kit v2 (Illumina, CA) and sequenced on the Illumina NextSeq 500. The data obtained and analysis is described in the S1 appendix. The candidate genes were shortlisted by using the flanking markers of the mapped *Nec3* physical region. The read pile-up from the *Nec3* region candidate genes were aligned from WT Bowman and *nec3-γ1* mutant (Bowman background) and deletions were observed in the gene specific read pile-up on the CLC Genome Workbench 8.0.3. (S3 Table)

### Candidate nec3 Gene Allele Analysis

The candidate *nec3* gene *HvCYP71-A1* was amplified from the *nec3-γ1* (Bowman background), WT Bowman, *nec3.1* (Steptoe background), *nec3.m* (Steptoe background), WT Steptoe, *nec3.d* (Proctor background), WT Proctor, *nec3.e* (Villa background) and WT Villa using the primer pairs Nec3_p450_F1 and Nec3_p450_R1; Nec3_p450_F3 and Nec3_p450_R2; Nec3_p450_F4 and Nec3_p450_R4 that were designed to produce overlapping amplicons with the respective fragment sizes of 1075, 563 and 404 bp, respectively (S4 Table). The purified PCR amplicons were sequenced using Sanger sequencing (Genscript, NJ). The overlapping PCR amplicon sequences were aligned and comparative analysis with the WT sequences was used to identify mutations for each of the independent *nec3* mutants.

### Exogenous Supplementation of Serotonin and Tryptamine

In three sets of treatments six replicates each of *nec3* mutants and their respective wildtypes were fed serotonin 150Μm, Tryptamine 5mM and water at 40 ml/plant through roots starting at 5 days old seedlings for every alternate day for total 10 feedings. The plants were inoculated with *B. sorokiniana* isolate ND85F spores at 2000/ml on the 14 days old seedling by following previously described protocol and phenotyped on day 21^st^ after 7 days of disease infection.

## SUPPLEMENTARY TABLE CAPTIONS

**S1 Table.** List of markers used to develop the barley *nec3* genetic and physical map on the chromosome 6H. The marker names, their sequences and physical positions are based on the 2019 Whole Genome Assembly of barley cultivar Morex.

**S2 Table.** The list of high confidence candidate genes in the delimited *nec3* region of the barley genome. The gene names, physical positions, their presence on the Exome Capture array and annotation is based on the 2019 Whole Genome Assembly of barley cultivar Morex.

**S3 Table.** The list of high confidence candidate genes in the delimited *nec3* region of the barley genome that were not captured in the exome capture experiment. These genes were further analyzed for their expression during the spot blotch infection at 72 hours post inoculation. The gene names, and annotations are based on the 2019 Whole Genome Assembly of barley cultivar Morex. Their relative normalized expression is reported in the Bowman wild type and *nec3-γ1* transcriptome profile post *B. sorokiniana* infection, where no-significant change from control is denoted by NS.

**S4 Table.** The List of primers used for the barley *Nec3* gene.

## SUPPLEMENTARY FIGURE CAPTIONS

**S1 Figure.** Typical reactions of *Bipolaris sorokiniana* isolate ND85F culture filtrates on *nec3* mutants and wildtype (wt) parental lines. Infiltrations of secondary leaves of barley lines Bowman wt, *nec3-γ1*, *nec3*.d, *nec3*.e., Steptoe wt, *nec3.l*, and *nec3.m* (left to right) with *Bipolaris sorokiniana* isolate ND85F culture filtrates (CF). The treatments from top to bottom (indicated to the left) were culture filtrates (CF) with Fries media (FM); CF with MOPS buffer (M); CF with M and pronase (P); and the control containing FM, M and P. Infiltrations were performed at the two-leaf stage and documented 4 and 7 days after infiltration. Pictures shown were taken at 7 days post infiltration.

**S2 Figure.** The *nec3* phenotype was not induced by chitin infiltrations on the *nec3* mutants and their respective wildtype barley plants. (A.) The panel shows the reaction to the chitin 2μg/ml infiltrations on the *nec3* mutants from the left *nec3.d, nec3.γ1, nec3.e, nec3.l* and *nec3.m*, followed by the Bowman, Steptoe, Proctor, Villa wildtypes. (B.) The panel shows the reaction to the control buffer infiltrations on the *nec3* mutants from the left *nec3.d, nec3.γ1, nec3.e, nec3.l* and *nec3.m*, followed by the Bowman, Steptoe, Proctor, Villa wildtypes.

**S3 Figure.** The *nec3* phenotype induced by *Bipolaris sorokiniana* inoculation on the secondary leaf of the *nec3* mutants (A.) and their respective wildtype barley plants (B.), where plants were supplied with 40 ml/pot of water; serotonin 150 μg/ml and Tryptamine 5mM on every alternate days starting from 5 day old seedling for a total of 10 root feedings.

**S4 Figure.** The mean lesion area observed by *Bipolaris sorokiniana* inoculation on the secondary leaf of the *nec3* mutants and their respective wildtype barley plants, where plants were supplied with 40 ml/pot of water; serotonin 150 μg/ml and Tryptamine 5mM on every alternate days starting from 5 days old seedling for a total of 10 root feedings. The lesions area were analyzed using ImageJ software for three independent leaves and 8 lesions per leave with the mean plotted for individual treatment along with the respective error bars.

**S5 Figure.** The transcriptome profile of the Bowman wildtype and nec3-γ1 mutant after 72 hours post inoculation with *Bipolaris sorokiniana* isolate ND85F. The venn-diagram represents the number of unique genes in a given class, where the blue and green oval represents Bowman downregulated and upregulated genes, respectively and the red and yellow oval represents *nec3*-γ1 mutant upregulated and downregulated genes, respectively (Top image). The bar graph shows the number of total genes present in each class of Bowman and *nec3*-γ1 mutant up and down-regulated genes (Bottom image).

**S6 Figure.** The NCBI blast phylogenetic tree of the barley Nec3 (unnamed protein product) protein, where the branch represents the relationship between the close orthologs of the proteins in monocots.

**S7 Figure.** The similarity between the barley Nec3 (HvCyp71A1_Nec3) and the rice SL (C71P1_ORYSJ) protein, where the dots represent same and the letters represents the different amino acid differences in the two proteins.

